# PhenotypeXpression: sub-classification of disease states using public gene expression data and literature

**DOI:** 10.1101/461301

**Authors:** Lucy Lu Wang, Huaiying Lin, Xiaojun Bao, Subhajit Sengupta, Ben Busby, Robert R Butler

## Abstract

The success of personalized medicine relies on proper disease classification and subtyping. Differential gene expression among disease subtypes can have a significant impact on treatment effect. This complicates the role of clinicians seeking more tailored diagnoses in cases where granular disease subtypes are not well defined. PhenotypeXpression (PhenoX) is a tool for rapid disease subtyping using publicly available gene expression data and literature. PhenoX aggregates and clusters gene expression data to determine potential disease subtypes, and develops a phenotypic profile for each subtype using term co-occurrences in published literature. Although the availability of public gene expression data is limited, we are able to observe clearly defined subtypes for several conditions.

## Introduction

Personalized medicine promises an almost limitless subdivision of clinical phenotypes. Accordingly, treatments can be tailored to individuals based on the specifics of their disease presentation (Mirnezami et al, 2012). In an effort to define and organize the rapid expansion of phenotypes and related conditions, a number of ontological systems have been developed, including the Online Mendelian Inheritance in Man (OMIM) (Hamosh et al, 2002), Orphanet (Weinreich et al, 2008), Human Phenotype Ontology (HPO) (Robinson et al, 2010), Human Disease Ontology (DOID) (Schriml et al, 2012), MedGen (Landrum et al, 2014), and others. These resources attempt to reach a consensus hierarchy for disease and a shared nomenclature. While unique identifiers are useful for resolving semantic differences, they do not provide an adequate description of the phenotype itself.

There has also been a steady increase in the amount of publicly available genetic expression data as public resources have scaled. The NCBI’s Gene Expression Omnibus (GEO) collects over 4000 gene expression datasets (Edgar et al, 2002). Several additional resources have been developed for expression data, such as the Human Cell Atlas (Regev et al, 2017),the Gene Expression Atlas (Kapushesky et al, 2010), the Cancer Genome Atlas (Weinstein et al, 2013), GTEx (Lonsdale et al, 2013), and others. These resources provide promising opportunities to aggregate and share expression data across multiple projects and for identifying novel phenotypes.

PhenotypeXpression (PhenoX) is a proof of principle tool for clinicians to quickly discover potential subtypes of known diseases. PhenoX rapidly aggregates gene expression data from multiple public studies, and then mines PubMed literature to develop novel disease subclassifications and expression profiles. By combining literature occurrences of subtype HPO and DOID information with GEO dataset expression profiles, each sub-classification of a disease is given a characteristic terminology fingerprint. Here we demonstrate the use of this tool for a common disease—Psoriasis—and identify two clusters with unique gene expression profiles.

## Software Description

### Operation

PhenoX synthesizes information from several resources: GEO, MeSH, HPO, and DOID. GEO, as an NCBI database, supports labeling of datasets and search using MeSH terms. Given an input disease term, PhenoX will identify the matches in the MeSH disease hierarchy and suggest near matches using string similarity. PhenoX also identifies any known and named subclasses of the disease from the MeSH hierarchy.

GEO profiles matching the MeSH term are filtered using the “up down genes” filter. Resulting matches are parsed to return a list of differentially expressed gene names and corresponding GEO DataSets (GDS) (Barrett et al, 2013). All GDS are retrieved and hierarchically clustered using R-pvclust hclust (Suzuki et al, 2006); the presence or absence of each gene in a GDS is represented as a binary variable and used for clustering. GDS with only a single differential gene are excluded to reduce noise. Clustering is based on euclidean distance in order to generate p-values via an approximately unbiased method with bootstrap probabilities at each node. Clusters are defined when the derived p-value is above the threshold of alpha greater than 0.95. After cluster assignment, the GDS in each sub-classification cluster is used for PubMed literature retrieval. To check for batch effects during clustering, the number of samples, date of submission, and platform type for each GDS is collected. Each cluster is compared to the overall distribution for a significant difference (α < 0.01) using the Kolmogorov-Smirnov test and chi-squared test for numerical and categorical data, respectively. Batch statistics are output to text file.

The PubMed identifier cited by each GDS is retained. Differential genes conserved across entire clusters are expanded using Hugo Gene Nomenclature (HGNC) gene names and aliases (Povey et al, 2001). This expanded gene list is used with the original MeSH term to query the PubMed database for additional PubMed articles potentially related to each cluster. For each cluster, we consolidate the titles, abstracts, and keywords corresponding to associated PubMed articles, and identify HPO and DOID terms within the article texts. The Spacy Lookup extension library is used for named entity recognition (NER) based on HPO and DOID dictionary terms. Term frequency data is gathered for each article, and the counts are merged for each cluster. Word clouds are generated and used to visualize the term frequency differences between clusters, with a separate term frequency list exported as a text file.

### Implementation

PhenoX is implemented in Python 3.6 and R. The minimum requirements, installation instructions, and usage instructions and examples are described in full on the PhenoX GitHub page. All Entrez data (GEO, GDS and PubMed) are accessed via Biopython (Cock et al, 2009). The software is designed to run on a personal computer, without requiring extensive resources. Usage is via the command line. The outputs include: (1) a dendrogram of matching GDS, (2) a word frequency visualization, (3) a heatmap of differentially expressed genes per GDS, (4) a pairwise distance graph of each GDS by absolute difference in gene sets, (5) a newick tree file, (6) a cluster batch statistics text file, and (7) a cluster term frequency text file.

## Results

### Computational Performance & Use Case

Using several MeSH terms as inputs (such as Psoriasis and Sepsis), the typical run time is 1-2 hours on a personal laptop with a 2.8 GHz dual-core processor and 8GB of RAM. The execution time is primarily split between PubMed article retrieval and NER, the latter occupying the majority of time. Clustering, including bootstrapping, takes less than ten minutes. Some disease terms may lack representative GDS (at least two required), and clustering cannot proceed.

In the case of Psoriasis (Figure 1), Cluster 1 contains a number of unique epidermal inflammatory or infectious diseases, including oral candidiasis, scabies, mucositis, and neurodermatitis. This suggests the possibility of comorbidities or confounders affecting some GDS groups (See Supplementary Table 1 for the full table of terms). In both the heatmap and the pairwise distance graph, there is consistent separation of two sub-groups, suggesting truly distinct expression profiles. Future work may include looking at deep phenotypes distinct in these two subgroups, and also looking at eQTLs in these subpopulations.

**Figure 1.**
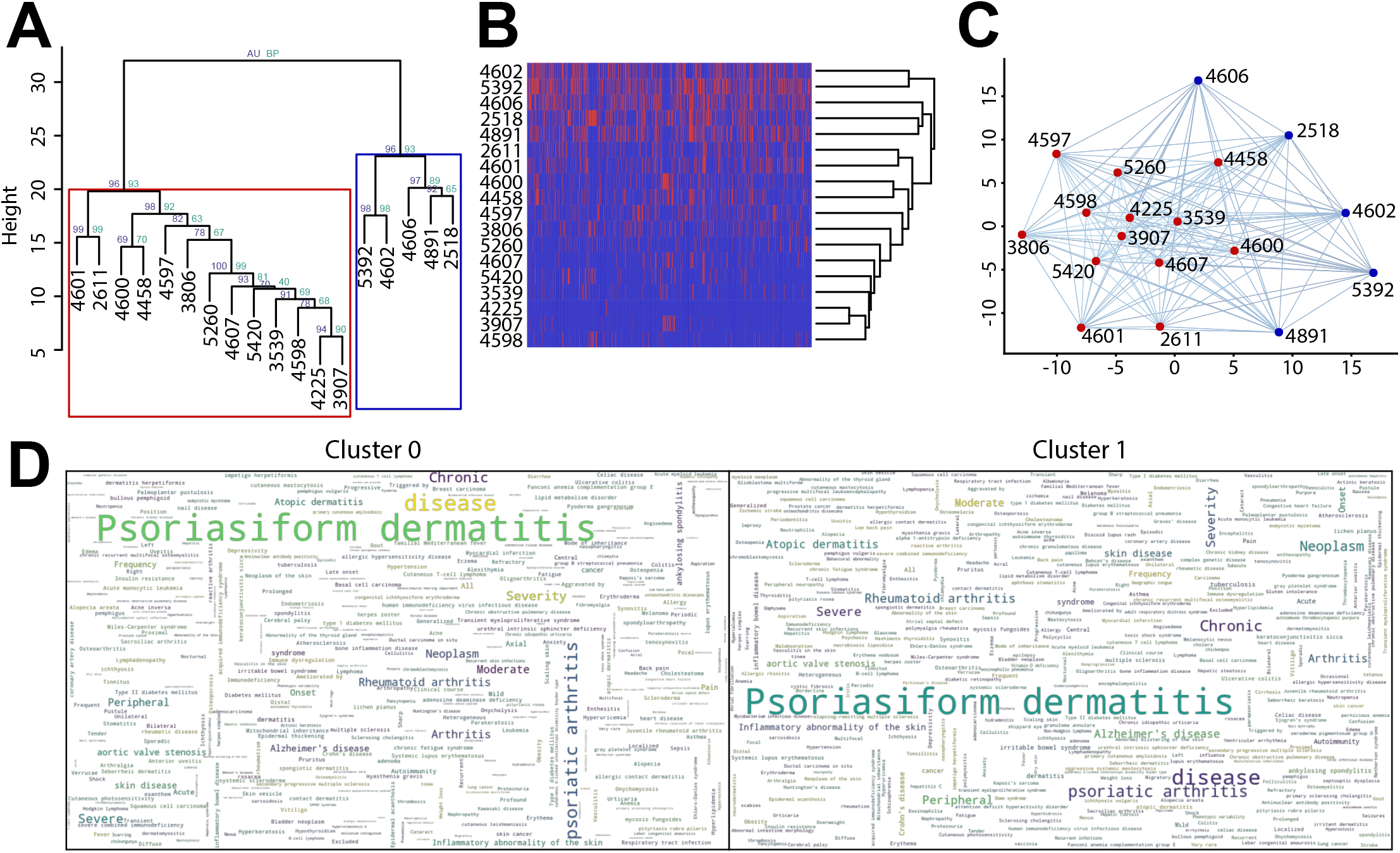
Visual outputs of PhenoX software. **(A)** A hierarchical clustering of GEO DataSets (GDS) using approximately unbiased (AU) and bootstrap probabilities (BP) with 5000 bootstraps. Clusters called at AU alpha > 0.95. **(B)** A differential gene expression heatmap depicting all available gene names associated with the searched term, with the presence indicated in blue and the absence in red. **(C)** A pairwise distance graph of each GDS by absolute difference in gene sets. GDS from Cluster 0 are colored in red, and those from Cluster 1 are colored blue. **(D)** A word cloud of DOID and HPO terminology associated with each cluster.

### Future Directions

These subtypes and terminology are exploratory results, inviting a number of hypotheses, such as: using subtype-associated genes to inform cohort selection, exploring treatment options by identifying associated gene pathways, and identifying potential comorbidities or confounders. Improvements to PhenoX’s interface could allow for more flexible data exploration. Introducing parallel processing in several steps, particularly NER, could also greatly shorten runtime. As PhenoX utilizes web services and produces relatively small output files, this software is ideal for a web-hosted graphical interface for researchers without computational backgrounds.

A significant limitation on PhenoX is the availability of curated expression data in GEO. The “up down genes” filter is only applied to a fraction of array data available in GEO (personal correspondence with GEO staff). According to the GEO website, “The subset effect scoring method is ad hoc…[t]his flag is simply an attempt to give potentially differentially-regulated genes higher visibility, and is not intended to provide an absolute determination of significance” (“Querying GEO”, 2016). A more thorough approach of analyzing each GDS to determine differential expression is beyond the scope of this project.

As a proof of concept, PhenoX is able to demonstrate a clear mechanism to subclassify disease populations based on gene expression, and to use that information to mine publically available literature. PhenoX is also amenable to future expansions of GEO expression cataloging, and can be modified to accept gene expression data from a growing number of resources. This includes technologies like RNA-seq that are easier to analyze in a programmatic fashion while correcting for batch effects.

## Data and software availability

Latest PhenoX source code available from: https://github.com/NCBI-Hackathons/phenotypeXpression.

Archived source code at the time of publication: https://doi.org/10.5281/zenodo.1479957.

A Docker image of PhenoX is available at:https://hub.docker.com/r/ncbihackathons/phenotypexpression.

Software license: MIT License

No competing interests were disclosed.

## Grant information

This research was supported by the Intramural Research Program of the National Library of Medicine, National Institutes of Health.

## Acknowledgements

We would like to thank Northwestern Feinberg School of Medicine for their support of the NCBI Hackathon where this work was created. We would like to thank Amazon Web Services for providing cloud computing resources for the event. We would also like to thank Paul Reyfman for organizing the event, and Karen Gutzman and Brett Cimbalik for librarian support.

